# Detection of *Campylobacter* in air samples from poultry houses using shot-gun metagenomics – a pilot study

**DOI:** 10.1101/2021.05.17.444449

**Authors:** Thomas H.A. Haverkamp, Bjørn Spilsberg, Gro S. Johannessen, Mona Torp, Camilla Sekse

## Abstract

**Background:** Foodborne pathogens such as *Campylobacter jejuni* are responsible for a large fraction of the gastrointestinal infections worldwide associated with poultry meat. *Campylobacter spp*. can be found in the chicken fecal microbiome and can contaminate poultry meat during the slaughter process. The current standard methods to detect these pathogens at poultry farms use fecal dropping or boot swaps in combination with cultivation / PCR. In this study, we have used air filters in combination with shotgun metagenomics for the detection of *Campylobacter* in poultry houses and MOCK communities to test the applicability of this approach for the detection of foodborne pathogens.

**Results:** The spiked MOCK communities showed that we could detect as little as 200 CFU *Campylobacter* per sample using our protocols. Since we were interested in detecting *Campylobacter*, a DNA extraction protocol for Gram negative bacteria was chosen, and as expected, we found that the DNA extraction protocol created a substantial bias affecting the community composition of the MOCK communities. It can be expected that the same bias is present for poultry house samples analyzed. We observed significant amounts of *Campylobacter* on the air filters using both real-time PCR as well as shotgun metagenomics, irrespective of the amount of spiked in *Campylobacter* cells, suggesting that the flocks in both houses harboured *Campylobacter* spp.. Interestingly, in both houses we find diverse microbial communities present in the indoor air. In addition, have we tested the *Campylobacter* detection rate using shotgun metagenomics by spiking with different levels of *C. jejuni* cells in both the mock and the house samples. This showed that even with limited sequencing *Campylobacter* is detectable in samples with low abundance.

**Conclusions:** These results show that air sampling of poultry houses in combination with shotgun metagenomics can detect and identify *Campylobacter* spp. present at low levels. This is important since early detection of *Campylobacter* in food production can help to decrease the number of food-borne infections.

## Background

*Campylobacter* spp. infection is one of the most frequently reported gastrointestinal infections of bacterial origin in Europe and worldwide [1, 2]. They cause campylobacteriosis with symptoms ranging from mild gastroenteritis to severe diarrhea. Complications can lead to a variety of diseases such as inflammatory bowel disease, reactive arthritis and Guillain-Barré syndrome [2]. For the genus *Campylobacter* more than 30 species and subspecies from several sources have been described and for many of these species the role as a human pathogenic bacteria is unclear [3]. Thermotolerant *Campylobacter* species, such as *C. jejuni* and *C. coli*, are most commonly associated with human infection and are often isolated from poultry and poultry products [1, 4]. *Campylobacter* has been isolated from the environment and a range of wild and domesticated animals, but poultry, especially broilers and laying hens, is considered as the main reservoir [5, 6]. In the EU, monitoring of *Campylobacter* is mandatory [7], and from 2018 monitoring of *Campylobacter* in broiler carcasses after chilling has been implemented in the member states [8]. In addition, to ensure a whole chain approach as recommended by EFSA, control measures should also be implemented at the farm level [9]. At present, on-farm sampling of poultry is carried out by sampling faecal droppings or using boot swabs, which are also widely used for *Salmonella* monitoring [10]. Interestingly, after *Campylobacter* colonize a flock, it is not only detected in faecal droppings and the litter, but also in the air inside the house. This knowledge was recently used to show that air sampling can be used as an alternative strategy for screening of *Campylobacter* in broiler flocks [11, 12]. Air sampling was demonstrated to detect the presence of *Campylobacter* and in some cases even earlier than the current conventional methods [11, 13, 14].

The air filters used, collect airborne material on a gelatin matrix, which is a product obtained from bovine or porcine skins and bones. The extraction process includes the use of extreme temperatures, pH and drying, which should create a sterile product. However, literature on contaminated gelatin indicates that common contaminants belong to thermotolerant, aerobic, endospore-forming bacteria. For instance, varieties of *Bacillus* species might be present, that are more resistant to the processes used in gelatin production and can therefore survive [15]. Thus, gelatin membrane air filters are sterilized using gamma irradiation before use, which should kill all organisms present [13]. Nonetheless, DNA of dead bacteria is likely still present in the gelatin matrix of the air filter.

In many air filter experiments, the collected airborne material is used for cultivation or specific PCR-based methods such as denaturing gradient gel electrophoresis (DGGE) or real-time PCR [11, 13, 16, 17]. The efficiency of cultivation is however suboptimal for specific organisms. A study by Johannessen et al., indicated that cultivation of air filters often failed to detect *Campylobacter* spp. as compared to boot swabs [11]. A follow-up study by Hoorfar et al supported this observation. This is in contrast to a higher *Campylobacter* detection rate when real-time PCR is used directly on the air filters compared to the same assay used material collected from boot swabs [11, 12]. The reason for the latter difference might be that the material on air filter is “cleaner” and therefore contain less inhibitors affecting PCR reactions.

Air filtering can collect all the microorganisms present in the enclosed air of buildings, which makes it suitable for diversity studies applying either amplicon or shotgun sequencing. Several studies describe poultry farm air samples indicating presence of a diverse community present in the poultry house air. For instance, Dai et al., showed that a range of potentially pathogenic bacterial genera, such as *Corynebacterium, Escherichia* and *Staphylococcus* are present in Chinese poultry house air [9]. Most of these organisms are associated to particulate matter particles present in the air with a particle size diameter smaller than 2.5 µm fraction [18]. Other studies using 16s rRNA DGGE profiling [16], shotgun sequencing [19], or cultivation [17] from material collected on air filters showed similar organisms being present in Canadian or Polish poultry houses. All these studies indicate a dominant airborne community dominated by Firmicute and Actinobacterial species, which are both phyla containing sporeformers [19]. In addition, it is also shown that these communities have a higher abundance of antibiotic resistance genes than found in the fecal material communities from which the farm air dust is originating [19]. As such, the presence of potential pathogens in the air of poultry houses therefore possess a possible health risk for both animals and workers.

The chicken fecal material deposited on the farm floor is partly responsible, via aerosol formation, for the bacterial community present in the poultry farm air. Therefor the chicken fecal microbiome has an influence on the air microbiome. The chicken fecal microbiome consist of a variety of species belonging to the Actinobacteria, Bacteroidetes, Firmicutes and Proteobacteria [20–23]. At a finer taxonomic scale, the dominant taxa are *Lactobacillaceae, Ruminicoccaceae* and *Enterobacteriaceae* in broiler fecal samples [23–25].

Pathogenic bacterial species such as *Acinetobacter, Campylobacter, Listeria, Proteus* and *Salmonella*, seem only present at low abundances in microbiomes from fecal material of healthy broiler flocks or are not present [23]. Interestingly, those taxa can be present on shells of eggs from the same animals that had a negative fecal sample [23]. Although fecal fluid can contaminate the eggshell, it is also possible that other sources, such as animal feathers, parental care, or even the environment could be responsible for the presence of pathogenic microbes. As such, sampling of fecal material might not give an adequate view of the presence of pathogens.

Here we tested the application of shotgun metagenomics on air filter samples spiked with mock communities and different numbers of *C. jejuni* to study the usefulness of the specific DNA extraction protocol as well as sequence depth in this study. In addition, air filter samples from two Norwegian poultry houses were tested as a proof of concept.

## Results

Four gelatin air filters (“MOCK” samples) were split into halves and spiked with either 75 or 125 µl of a mock community consisting of cells from 10 different species in a log abundance distribution (Figure 1, Table 1). Two of the half filters spiked with mock community were further spiked with 200 CFU (colony forming units) of *C. jejuni* CCUG 11284T and another two with 20.000 CFU, while the remaining four halves were not spiked and used as negative controls. One air sample from each of the two poultry houses on the same farm were collected (“HOUSE”). Both filters were split in four pieces. One piece from each filter was not spiked, while the other three were spiked with three different levels of *C. jejuni* 927 (Figure 1, Table 1). All filter pieces were treated as separate samples.

**Table 1.** Description of samples and *Campylobacter* 16S rDNA real-time PCR results. Samples are listed by levels of spiking.

| Sample ID | Sample -<br>*C. jejuni* spike | Spike<br>(CFU) | Mock amount<br>(µl) | *Campylobacter*,<br>Cq |
| --- | --- | --- | --- | --- |
| 5-MOCK1-S2a | Mock - CCUG 11284T | 20000 | 75 | 29.1 |
| 6-MOCK1-S2b | Mock - CCUG 11284T | 20000 | 75 | 29.2 |
| 3-MOCK1-S1a | Mock - CCUG 11284T | 200 | 75 | 37.2 |
| 4-MOCK1-S1b | Mock - CCUG 11284T | 200 | 75 | 35.2 |
| 1-MOCK1-Ba | Mock – no spike | 0 | 75 | 43.1 |
| 2-MOCK1-Bb | Mock – no spike | 0 | 75 | No Cq |
| 7-MOCK2-Ba | Mock – no spike | 0 | 125 | 41.1 |
| 8-MOCK2-Bb | Mock – no spike | 0 | 125 | No Cq |
| 9-H1-3 | House 1 - 927 | 5650 | - | 29.4 |
| 10-H1-4 | House 1 - 927 | 565 | - | 30.4 |
| 11-H1-5 | House 1 - 927 | 56,5 | - | 30.3 |
| 12-H1-B | House 1 – no spike | 0 | - | 31.1 |
| 13-H2-3 | House 2 - 927 | 5650 | - | 30.6 |
| 14-H2-4 | House 2 - 927 | 565 | - | 31.3 |
| 15-H2-5 | House 2 - 927 | 56,5 | - | 31.3 |
| 16-H2-B | House 2 – no spike | 0 | - | 32.6 |

**Figure 1.**
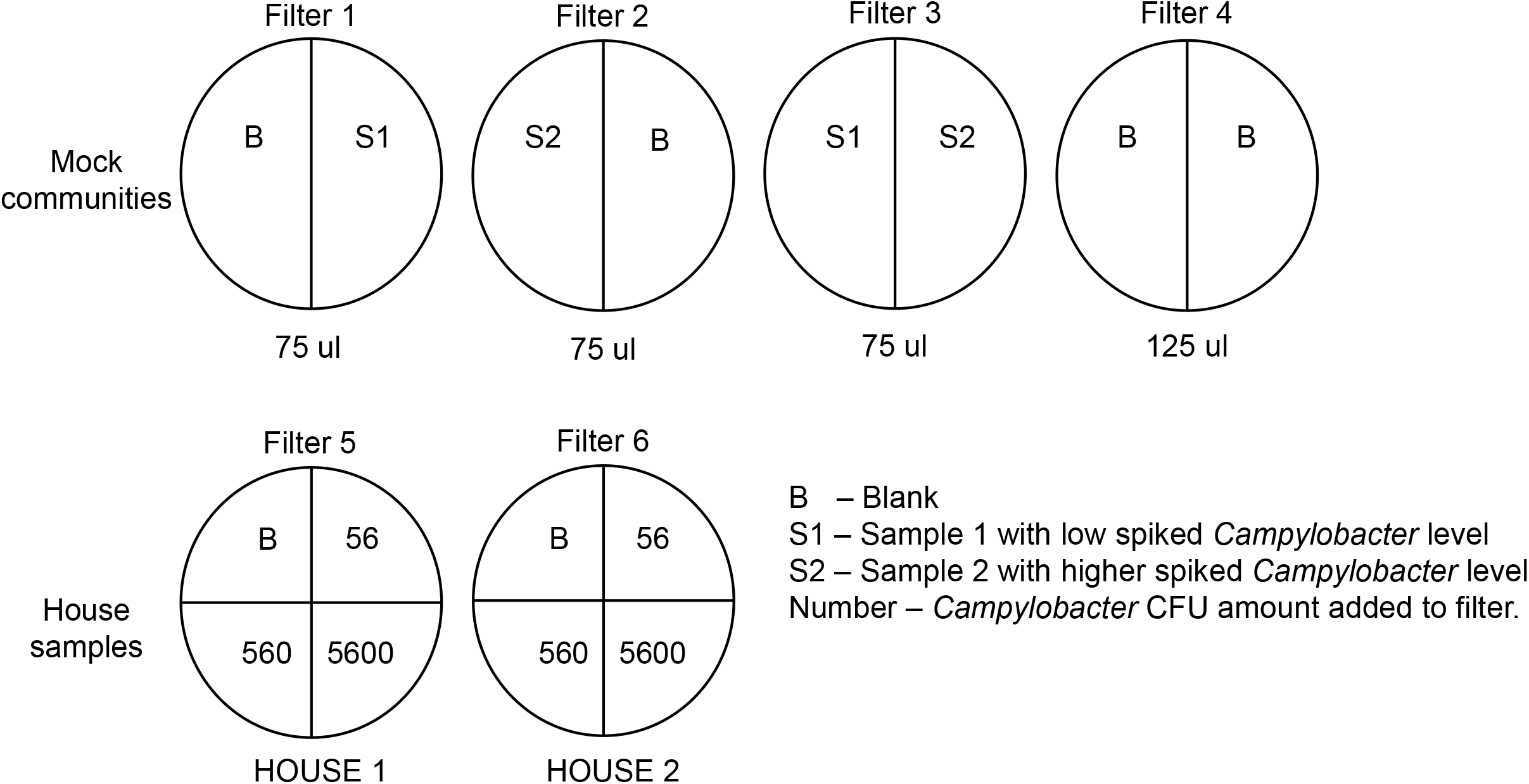
Design of the experiment using different air filters. The MOCK samples were spiked with 200 (S1) or 20.000 (S2) CFU of *C. jejuni* CCUG 11284T. The HOUSE samples were spiked with three different levels of *C. jejuni* 927.

Real-time PCR was used to detect *Campylobacter* in all samples. Campylobacter was detected at approximately Cq 29 in the samples spiked with 20.000 CFU and at approximately 36 in the samples spiked with 200 CFU (Table 1). The MOCK samples that were not spiked with *Campylobacter* were all negative (two “No Cq” and two above our standard hard cutoff Cq value of 40, Table 1). *Campylobacter* was detected in all HOUSE samples including the non-spiked samples, demonstrating that both poultry houses harboured *Campylobacter*.

### Microbial community composition

Microbial community composition of the air filter communities was determined using metagenomic shotgun sequencing (Table 2). In order to assess which fraction of the microbial communities was captured using shotgun sequencing we used Nonpareil3 [26]. This indicated that for the MOCK community samples we had reached sufficient coverage (> 0.97), while the HOUSE samples with a chicken house background had an average coverage of 0.76. This result was expected as the MOCK communities only consisted of 10 species, while the HOUSE communities were natural samples with many more. Thus, in line with this, we find that the HOUSE samples showed a higher Nonpareil index of sequence diversity (N_d_) (19.3 ± 0.1) than the MOCK samples (17.6 ± 0.2) [26]. This index describes the community diversity in sequence space using a natural logarithm. The N_d_ found for our MOCK samples falls in the same range as Mock communities tested by the authors of Nonpareil3. Most host-associated communities tested in Rodriquez-R et al., show a N_d_ range between 17 and 22, and our HOUSE samples show similar N_d_ values [26].

**Table 2.** Overview of the sequencing effort for each of the samples in this study.

| Sample ID | *C. jejuni*<br>cfu | Raw<br>Paired-<br>end<br>sequences | After quality<br>trimming<br>/filtering (%) | PhiX<br>sequences | Human<br>sequences | Final PE<br>sequences | Metagenome<br>Coverage | Sequence<br>diversity<br>index (Nd) | Average<br>genome size<br>(Mbp) |
| --- | --- | --- | --- | --- | --- | --- | --- | --- | --- |
| 1-MOCK1-B-a |  | 24675209 | 93.8 | 12 | 16022 | 21444439 | 0.99 | 16.0 | 4.10 |
| 2-MOCK1-B-b |  | 18611989 | 92.6 | 11 | 6553 | 16267510 | 0.98 | 16.1 | 3.80 |
| 3-MOCK1-S1-a | 200 | 21636112 | 93.2 | 16 | 6197 | 18775078 | 0.98 | 16.1 | 3.95 |
| 4-MOCK1-S1-b | 200 | 25799040 | 93.7 | 18 | 44918 | 22326211 | 0.99 | 15.9 | 4.30 |
| 5-MOCK1-S2-a | 20000 | 25760599 | 92.3 | 16 | 33337 | 22067796 | 0.98 | 16.0 | 4.02 |
| 6-MOCK1-S2-b | 20000 | 19029552 | 93.9 | 17 | 16598 | 16649467 | 0.98 | 16.0 | 4.08 |
| 7-MOCK2-B-a |  | 16256676 | 92.7 | 25 | 52135 | 14515655 | 0.99 | 15.3 | 3.38 |
| 8-MOCK2-B-b |  | 17247065 | 90.9 | 11 | 8124 | 14417819 | 0.99 | 15.9 | 4.21 |
| 9-H1-3 | 5650 | 16183867 | 96.4 | 10 | 4236 | 14063204 | 0.74 | 19.4 | 3.24 |
| 10-H1-4 | 565 | 15842872 | 96.2 | 7 | 3936 | 13728192 | 0.74 | 19.3 | 3.20 |
| 11-H1-5 | 56 | 26674096 | 96.2 | 15 | 7839 | 23171336 | 0.78 | 19.4 | 3.20 |
| 12-H1-B |  | 31645569 | 97.0 | 10 | 9333 | 27869436 | 0.79 | 19.4 | 3.24 |
| 13-H2-3 | 5650 | 23296681 | 96.1 | 14 | 20021 | 20297450 | 0.79 | 19.2 | 3.14 |
| 14-H2-4 | 565 | 20578293 | 95.9 | 15 | 4504 | 17927138 | 0.78 | 19.2 | 3.12 |
| 15-H2-5 | 56 | 13873052 | 95.9 | 6 | 2728 | 12152054 | 0.75 | 19.2 | 3.14 |
| 16-H2-B |  | 15071970 | 96.2 | 11 | 3183 | 13215124 | 0.76 | 19.2 | 3.19 |

The taxonomic community composition of the air filter communities was analyzed using Kraken [27]. For the MOCK communities we found that on average 9 % of the reads were unclassified, while for the HOUSE communities this was 48.8 %. At the phylum level we detect 10 taxa with a relative abundance >= 0.01 % in most of the samples (Supplementary materials figure S1). Firmicutes and Proteobacteria dominated the MOCK samples, while in the HOUSE samples we found the same together with the phyla Actinobacteria and Bacteroidetes. The other eight phyla were all present in the HOUSE samples at relative abundances below 0.02 % except for Deinococcus-Thermus (average relative abundance = 0.023 ± 0.003%). The later thermophilic phylum was present in all our samples and was likely a contaminant from the air filter gelatin matrix.

The main bacterial genera found in the MOCK samples were *Listeria* and *Pseudomonas* and they showed relative abundances deviating from the expected abundance (Figure 2). Both taxa were predicted to have abundances of 89.9% and 8.9%, but instead show relative abundances of 37.9% (±13.5) vs 42.7% (±14.8). This indicates that our DNA extraction protocol, aimed to extract DNA from Gram-negative bacteria, caused an underestimation of Gram-positive bacteria in our samples.

**Figure 2.**
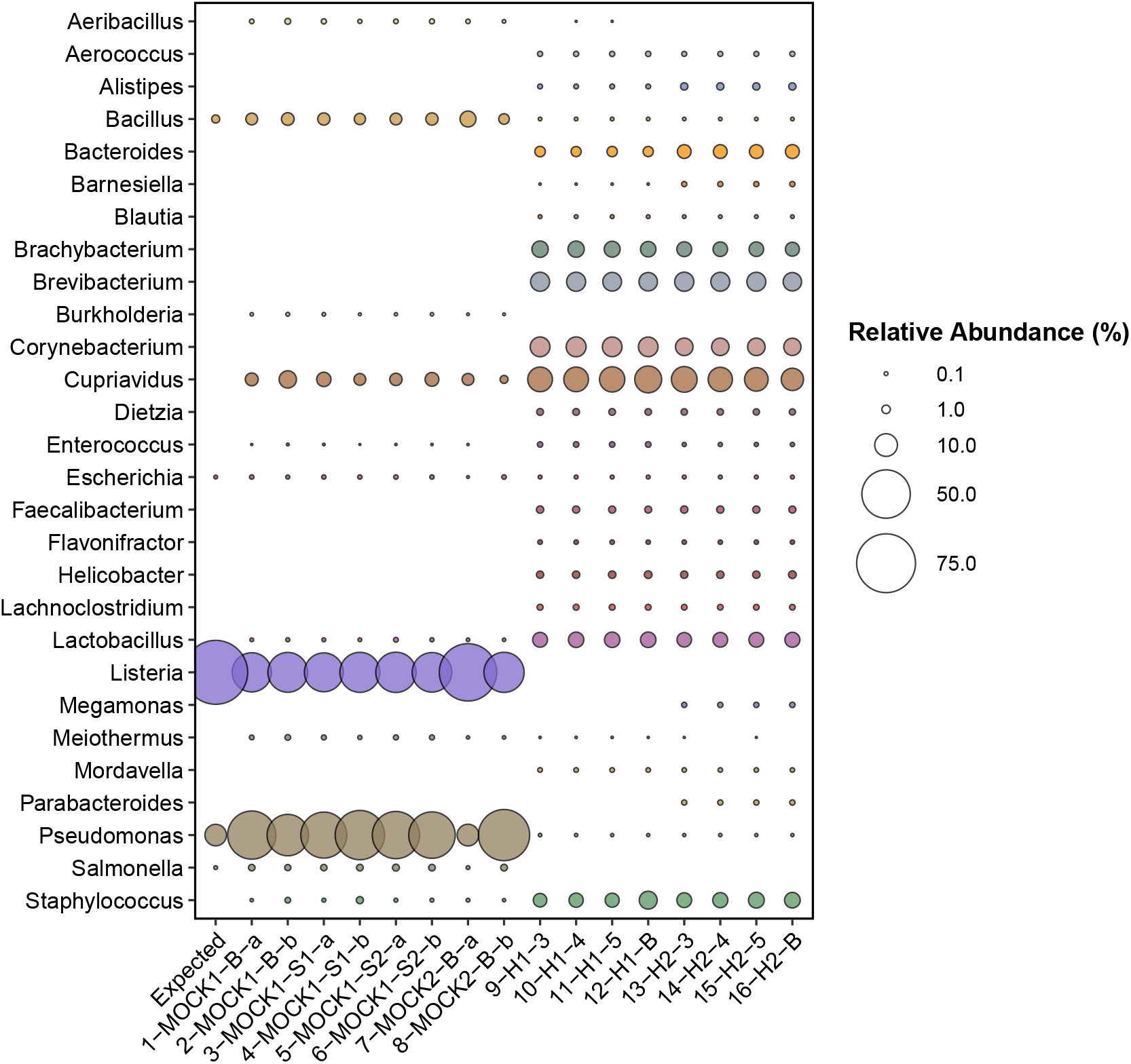
Relative abundances of bacterial genera found in MOCK and HOUSE samples. The first column shows the expected abundances of MOCK community taxa as provided by the manufacturer. Genera with a relative abundance equal or higher than 0.1% in at least one sample were kept for visualization. Taxa relative abundances below 0.01% are not shown.

In the HOUSE samples we found a diverse group of different genera with some known to contain pathogenic species (Figure 2). The dominant taxa were *Brevibacterium, Brachybacterium, Bacteroides, Corynebacterium, Lactobacillus, Staphylococcus, Faecalibacterium*, and *Helicobacter*. Most taxa were found with similar abundances in both houses, but a few had higher abundances in the HOUSE 2 samples (Figure 2). Those taxa were *Bacteriodes, Alistipes* and *Megamonas*. Overall, there is a high reproducibility between the samples from a single house.

A few genera are especially notable, since they were detected in both HOUSE and MOCK samples (Figure 2). A more detailed analysis showed that all samples contain several thermophilic bacteria, e.g. *Aeribacillus, Meiothermus* and the Betaproteobacterial genus *Cupriavidus*, while the MOCK samples also contained the Betaproteobacterial genus *Burkholderia*. Since these genera are not part of the MOCK community, it is likely that they are contaminants present in the gelatin matrix of the air filter. Interestingly, *Cupriavidus* was present with more than 1 million reads (max 4 Million) in the HOUSE samples, while in the MOCK communities they only showed up with on average 0.5 million reads. Although the MOCK communities were created using only 10 distinct species, we detected many more species although most were in low abundance (< 0.01%). These classifications are likely due to misclassifications. For instance, in the MOCK communities that were not spiked with *Campylobacter*, we consistently detected this genus using Kraken with an average classified read count of 43 (Figure 3A). A similar number was identified when we mapped the metagenomics reads against a collection of *Campylobacter* genomes and the two genomes of our isolates used for spiking (Figure 3B). This indicates a consistent detection irrespective of the method used for detection. For the MOCK communities spiked with 200 or 20000 CFUs we could classify on average 100 or 6825 reads respectively. These results indicate that species detected using Kraken with close to 50 reads or less are likely not present in our samples, and should be ignored. Nonetheless, even with the restriction that a species has to be present in all MOCK samples with more than 100 reads, we still find 121 genera present.

**Figure 3.**
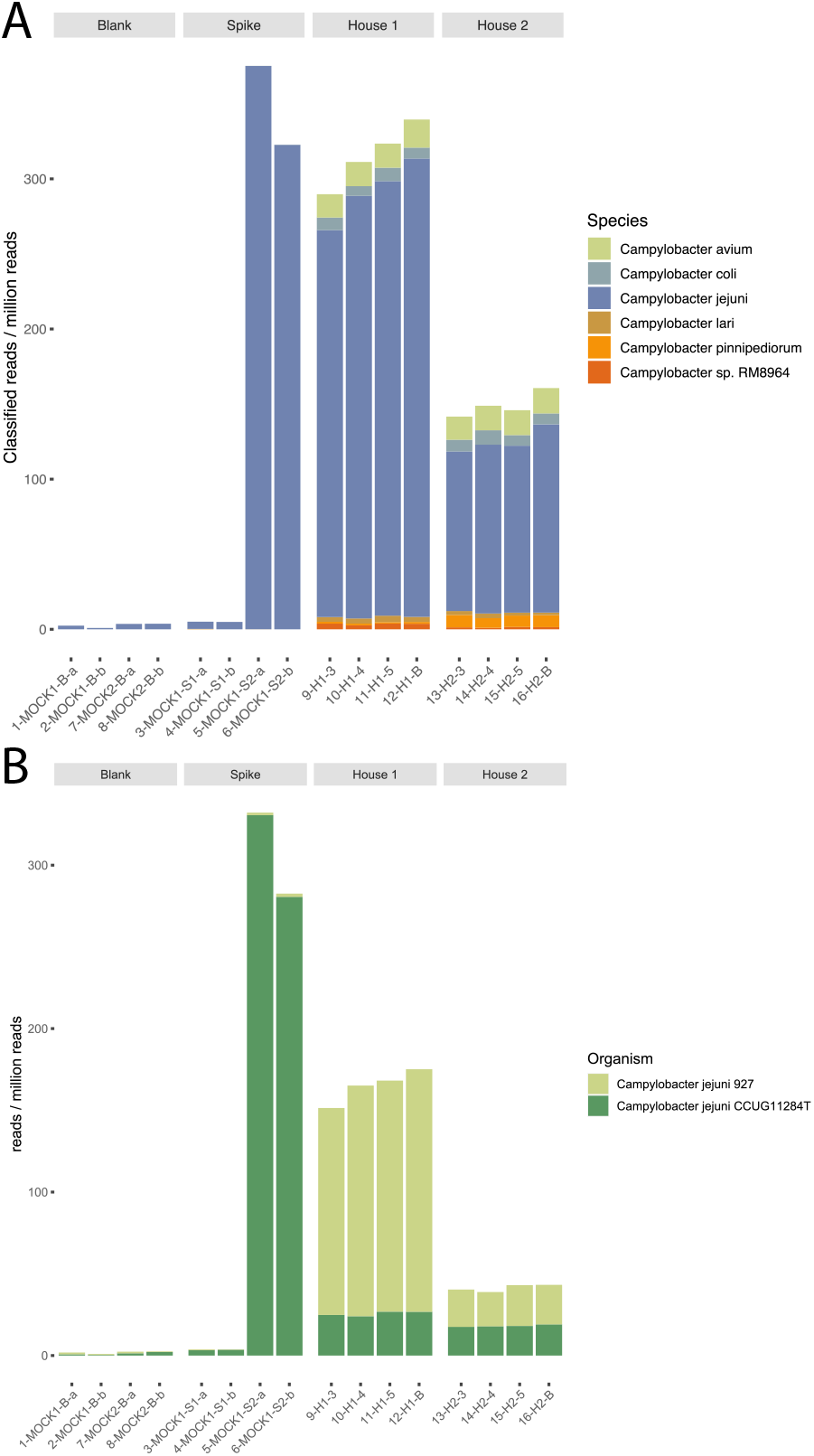
*Campylobacter* spp. abundances in MOCK and HOUSE samples using two different methods. **A**) Normalized abundances of *Campylobacter* spp. as found with Kraken read classification. Read abundances for the different taxa are shown as classified reads per million reads. **B**) Normalized abundances of reads mapped by BBsplit to the genome sequences of *C. jejuni* CCUG 11284T (MOCK sample spike) or *C. jejuni* 927 (HOUSE sample spike). Note that in the HOUSE samples we find reads mapping to *C. jejuni* CCUG 11284T, indicating the presence of strains closely related to this isolate.

### Campylobacter detection of communities spiked with C. jejuni

We used both Kraken taxonomic classification as well as mapping with BBsplit to isolate genomes to identify *Campylobacter* spiked into MOCK and HOUSE samples. Both methods identified reads matching to *C. jejuni*, but since the genomes were not available, neither Kraken not BBsplit could differentiate at the strain level for the two strains used for spiking. We thus sequenced the two isolates *C. jejuni* 927 and *C. jejuni* CCUG 11284T with Illumina HiSeq 3000 and used the resulting assemblies to map our metagenomics reads.

The draft assemblies for *C. jejuni* 927 and *C. jejuni* CCUG 11284T were submitted to ENA (accession numbers: CAJPVE01; CAJQFQ01) and have a genome size of 1.59 Mbp and 1.73 Mbp respectively (Supplementary table 1). We used the Tetra correlation search at the JspeciesWS webserver [28] to identify the taxonomically closest known isolates for our strains. These isolates are *C. jejuni* subsp. *jejuni* 327 for 927 (Average Nucleotide Identity (ANI): 100.00) and *C. jejuni* subsp. *jejuni* ATCC33560 for CCUG 11284T (ANI: 99.66). The match for CCUG 11284T indicates the same strain but with a different culture collection identifier. The ANI value between 927 and CCUG 11284T is 97.5%.

With the mapping approach, we find a clear difference between the samples on how abundant either spike isolate was (Figure 3B). In the MOCK samples that was spiked with CCUG 11284T we find reads uniquely matching C. *jejuni* 927 (Figure 3B). Likewise, reads with a unique match to CCUG 11284T were found in the HOUSE samples that were spiked with C. *jejuni* 927. In addition, do we find many more *Campylobacter* species in the HOUSE samples (Figure 3A, Supplemental figure S2B).

## Discussion

### Campylobacter detection using metagenomics

The prevention of gastrointestinal disease due to foodborne pathogens requires early and sensitive detection of pathogens along the food production chain. For pathogens such as *Campylobacter* spp., it is important to identify the presence at the farm to prevent (or limit) further contamination. The current method for on-farm sampling of poultry is by boot swabs of the floor environment, including fecal droppings, analyzed by either cultivation or real-time PCR. These methods may have limitations in their application due to cultivation bias or PCRs limitation to one target per assay. In recent years, shotgun metagenomics has been shown to be able to detect a large variety of microbes without cultivation and as such, it can be used to identify multiple pathogens simultaneously. Furthermore, air sampling at poultry farms combined with PCR detection indicated earlier detection of pathogens that conventional sampling [11, 12]. In addition, the DNA isolates from the air samples did contain less PCR inhibitors. Thus, combining metagenomics with air sampling could help to identify a variety of pathogens without being affected by cultivation biases.

Here, we used real-time PCR and metagenomic shotgun sequencing to analyze microbial communities present in MOCK and HOUSE samples with a specific focus on the detection of *Campylobacter*. With real-time PCR we were able to detect *Campylobacter* in the MOCK communities when spiked with two different levels (Table 1). For the HOUSE samples we found Cq values between 29 and 33, with the lower values in the non-spiked samples (Table 1). The detection of *Campylobacter* in the non-spiked sair filters demonstrated that both poultry houses were infected with *Campylobacter*. These results were also confirmed by cultivation of boot swabs sampled simultaneously as the air samples from the two houses [11].

With the shotgun data we used two different approaches to identify *Campylobacter* spp. in our samples (Figure 3). The classification approach with Kraken showed that the HOUSE samples had between 100 and 300 reads / million reads that could be classified to *Campylobacter spp*. Interestingly, BBsplit mapped between 40 and 170 reads / million reads to both genomes of the spiked isolates, which is a slightly smaller fraction of the metagenomics datasets. BBsplit was used with a with a variety of Campylobacter spp genome, and it only counts reads mapping unambiguously to a single genome. When a read would map equally well to multiple genomes it was not counted. Interestingly, the mapping showed that in the HOUSE samples strains *C. jejuni* CCUG 11284T and 927 were found. The latter was spiked into these samples, while the former was used in the MOCK communities as a spike. This suggests the presence of *C. jejuni* strains sharing genomic similarity with CCUG 11284T in the HOUSE samples. An additional 0.03 % of reads mapped to a variety of other *Campylobacter* spp. including a third *C. jejuni* (NCTC 11168) reference genome (Supplementary materials Figure S2). In contrast, we only find few reads (<1 read / million reads) in the MOCK samples that were assigned to *C. jejuni* 927, while 3 (200 CFU) to 300 reads / million reads (20.000 CFU) were assigned to *C. jejuni* CCUG 11284T in the spiked samples.

Our results suggest that both Kraken and BBsplit can be used to identify *Campylobacter spp*. in the air of Poultry houses as does real-time-PCR [11]. The added value of using shotgun metagenomics for detection is the possibility to identify a variety of *Campylobacter* species and other bacteria of interest present at a farm, which is not possible with single 16s rRNA based real-time PCR. However, our metagenomic results also indicate that interpretation of such data should be done with care. For instance, the BBsplit results suggests the presence of a large number of different species (supplementary materials Figure S2), while Kraken predominantly identified *C. jejuni* (Figure 3). Some of the species genomes used in the BBsplit mapping approach have very distinct host ranges. For instance, the species *C. fetus, C. hyointestinalis* and *C. iguaniorum* are associated with other hosts [29–31] and because of that it seems unlikely that they are present in our samples. Others like *C. jejuni* are known to be genomically versatile with broad ecological/ host ranges, including wild birds, livestock and humans [32, 33]. In addition, *Campylobacter* spp. *are* known for their high rate of horizontal gene transfer making it likely that environmental isolates from different species might contain genome regions shared by multiple species [34, 35]. That makes it difficult to identify these different species unambiguously and caution is needed for the interpretation. Nonetheless, by using two different approaches we show that multiple *Campylobacter* species are likely present in the air of our HOUSE samples and that most are at low abundance.

### Air sample microbiomes vs broiler fecal microbiomes

The microbiome composition of the air in poultry houses is related, but not similar, to the fecal microbiome composition. After deposition of the feces on the litter layer in the house the microbiome composition likely changes in community composition and only a part of the community becomes airborne. In our analysis, we find a highly diverse community present in the air, with several genera also known to be present in the broiler feces. Typical genera found in broiler feces are *Bacteriodes, Brevibacterium, Corynebacterium, Enterococcus, Lactobacillus and Staphylococcus* [22– 24, 36, 37]. These taxa were also identified in our air sample microbiomes of the poultry houses as well as in other studies using different detection methods [16–18]. That suggests that the air microbiome in poultry houses can be used as a proxy for the fecal microbiome. It also implies that the poultry house air contains many of the microbes associated with the animals and as such gives a good overview of the microbiome of the entire flock. Interestingly, many of the genera identified in our analysis are known gut microbiome species, but not all of them. *Brachybacterium* spp. for instance can be isolated from a wide range of sources including poultry litter, Gouda cheese, oil brine or via air sampling [38, 39]. Not unexpected, *Brachybacterium* spp. have also been identified in the dust from poultry farms [18]. They have been rarely identified as pathogenic agents in humans [40]. The same is true for the genera *Dietzia and Pseudomonas*, which are both widely distributed bacteria that can be opportunistic pathogens [41]. Interestingly, *Pseudomonas* spp. can regularly be found in chicken fecal material as well as the air of poultry houses and are of concern for food production [17, 23, 42, 43]. These results therefore indicate that shotgun metagenomics of air samples provide a suitable approach to monitor poultry farms for a wide variety of pathogens.

### Sensitivity of air sample metagenomics

The use of air sampling combined with real-time PCR or sequencing to detect pathogens at poultry farms has clear benefits over the gold standard with boot swabs with respect to the higher sensitivity and faster throughput [11, 12]. An additional advantage is that the potential to use the same samples to study all the microbial species present with metagenomic approaches. However, there is some ptifalls for these approaches. The first factor is the low input amount of DNA collected by this technique. Such samples are highly sensitive to contaminants from kits and reagents themselves, in addition to laboratory contamination [44, 45].

In addition, the DNA extraction method can introduce bias. Here, we used a method aimed at Gram-negative bacteria and specifically at lysing the cells from *Campylobacter* spp.. Using MOCK communities, as expected we find a large difference between the expected and observed MOCK community composition of the samples. This was especially clear for the abundances of *L. monocytogenes* and *P. aeruginosa*, which had a predicted relative abundance of 89 vs 8.9 %, but instead show an almost equal abundance in this experiment (Figure 2, supplementary figure S3). This was most likely due to the gram positive *L. monocytogenes* DNA being extracted with lower efficiency that the gram negative *P. aeruginosa*. Nonetheless, in the HOUSE samples, we do find many gram-positive bacterial genera, but the relative abundance of these taxa is likely underestimated relative to Gram-negative bacteria. This underscores the importance of including mock communities in microbiome studies, in order to understand the DNA extraction bias for particular communities.

Our results suggest that the gelatine filters contain DNA despite the fact that these filters are gamma sterilized [15]. Gelatine production is a harsh, but not a sterile process, which requires sterilization before shipment of such filters. By using mock communities we were able to identify several contaminating genera likely present in the gelatin matrix of the air filter. The taxa *Meiothermus, Aeribacillus* and *Burkholderia* could be found in all MOCK samples and with similar abundances in some of the HOUSE samples. More interestingly, we also find a high abundance of *Cupriavidus* in all samples, which strongly suggests this is a contaminant genus. This *Burkholderiales* genus, together with *Burkholderia* and the sister genus *Ralstonia* can be found as contaminants in microbiome studies, as shown previously [44, 46]. This genus has also been found in another air sampling study of poultry houses where polyvinyl chloride filters were used [16], but there is no indication that the authors of that study controlled for contamination by analyzing clean filters. Thus, it is unclear if this genus is part of the poultry house microbiome as indicated by Just et al [16], or that it is also present as a contaminant in their non-gelatine filters. Since we observe *Cupriavidus* in all our air filter samples it is unlikely that this genus is present in the poultry house, but rather introduced into the samples via the gelatin matrix or through laboratory handling. What is clear from our analysis is that contaminating species in air filter experiments are present and easily detected when using mock communities or even negative samples. This indicates a need to include such sample types in all microbiome experiments using air sampling, to be able to filter out contamination from the filters.

### Application of air sampling

Our study showed that air sampling indoor air of poultry houses together with shotgun metagenomic sequencing can be used to detect pathogenic microorganisms. This approach can be used to study a various of pathogens, moulds, antimicrobial resistence in environments where a high hygiene standard is required [19, 47]. These findings indicate that in l.e. animal production facilities, slaughter houses or in the food industry airborne pathogens and their resistome might cause health concerns for the humans working in such spaces [17, 48]. Likewise, the animals in these facilities are also exposed to the same potential pathogens that are present in the air [18, 49]. With the increasing availability of shotgun metagenomics, it becomes possible to monitor animal house facilities for a wide variety of pathogens without the limits in detection when using cultivation or dedicated PCR approaches. This enables the industry not only to monitor for pathogens that can cause foodborne outbreaks at an early stage, but it could also stimulate the development of intervention methods to reduce the proliferation of such pathogens at the farm level. In addition, these approaches can help to improve the animal health by developing measures to reduce the exposure to a wide variety of potential pathogens. Such measures could include better regulation of the indoor climate of poultry houses. The extra benefit of such developments is that the working conditions for humans at those farms might improve as well.

Nonetheless, to really make use of the power of metagenomics in an industrial monitoring set-up, it will be needed to further reduce the cost of this method, and to develop analysis approaches that are both user friendly and easy to use for non-bioinformaticians. Finding potential pathogens in the air of poultry farms might cause concern, but since many of the organisms found are also part of the healthy fecal microbiome of chicken it is difficult to identify at what level pathogen detection might require intervention. These difficulties require further investigations in order to improve the sensitivity and specificity of metagenomic detection methods.

## Conclusions

Here we used shotgun metagenomics and real-time PCR to study the microbial community present in the air of poultry houses. Our results indicate the presence of a diverse community that contains many genera that are also found in the chicken fecal microbiome. This shows that air sampling of poultry house air combined with shotgun metagenomics can be used as a proxy for the study of the fecal microbiome of the flock present in a farm house. The shotgun metagenomics results allowed us to identify *Campylobacter* spp. present at low levels, as well as other pathogenic microbes that could be of health concern. By using mock communities, we address some technical limitations that can help data analysis and interpretation of data generated with the air sampling method. Those include DNA extraction biases due to the extraction method chosen, as well as the background contamination of the samples due to laboratory handling and/or background DNA present in the gelatin matrix of the filters.

By using air sampling methods with shotgun metagenomics it will be possible to study the microbial communities associated with animal production and food industry with emphasis on the presence of microbes considered as food-borne pathogens. Understanding the dynamics of microbial communities present in poultry houses will help to identify processes to reduce to the introduction of pathogens in the poultry production as well to improve poultry and human worker health.

## Methods

### Sample collection

Air filters from two Norwegian broiler houses on the same farm were collected using an AirPort MD8 device (Sartorious Stedim Biotech, France) with disposable gelatin filters (80 mm diameter; Sartorius, 17528-80ACD) as described by Johannessen and coworkers [11], one from each house. The filters collected were used for spiking as described below.

### Preparation of artificially contaminated air filters

Four gelatin membrane filters were divided into two pieces each (Table 1, Figure 1). Six half filters were spiked with 75 µl of microbial community standard (MOCK), and two half filters were spiked with 125 µl of the same standard. The microbial community standard (MOCK) used in this study has a log abundance distribution of different bacterial species and fungi ranging from 10^8^ to 10^2^ cells (ZymoBIOMICS Microbial Community Standard II, ZYMO RESEARCH EUROPE GMBH, Freiburg, Germany) and is well suited for assessing the accuracy of composition and ideal for the quality control of microbiome measurements. In addition, we spiked four of the half filters already spiked with mock communities, with two different concentrations of *C. jejuni subsp. jejuni CCUG 11284T* (Accession number CAJQFQ01) (200 and 20.000 cfu) (Table 1). One air filter collected from each of two broiler houses (see above) were divided into quarters and spiked with three different concentrations of *C. jejuni* 927 (Accession number CAJPVE01) (5600, 560 and 56 cfu) (Table 1). One quarter of each filter was not inoculated with *C. jejuni* 927 and will reflect the content of the real sample (Figure 1).

The DNA extraction is performed as previously described in Hoorfar et al., 2020 [12]. Briefly, air filters were pretreated for DNA extraction as follows; half a filter was dissolved in 3.5 ml ddH_2_O and 100 µl alkaline protease (Protex 6L, Genencor International BA, The Netherlands) was added. The samples were mixed by vortexing until the filters were dissolved. The samples were then incubated at 37°C in a thermal mixer for 6 min at 1000 rpm, centrifuged at 4°C for 5 min at 8000 x g and the supernatant discarded. The pellet was used for DNA extraction using the Gram-negative protocol for DNeasy Blood and Tissue kit (Qiagen) with some minor modifications described briefly below. The incubation period at 56°C was set to one hour for all samples and all samples were treated with 4 µl RNase A (100 mg/ml, Qiagen). DNA was eluted in 100 µl TE buffer with 0.1 mM EDTA. The DNA quantity was determined by using the high sensitivity DNA Qubit assay (ThermoFisher Scientific) and DNA quality was assessed, if enough DNA, by using Nanodrop spectrophotometer (ThermoFisher Scientific).

DNA from the two *C. jejuni* strains used for spiking were extracted using QIAamp DNA Mini kit (Qiagen) according to the manufacturer’s protocol. DNA quantity and quality was determined by using the broad range DNA Qubit assay and Nanodrop spectrophotometer. The DNA from the two *C. jejuni* isolates was whole genome sequenced using the same protocols as for the metagenomic samples, for details see below under metagenomic library preparation and sequencing.

#### Real-time PCR for Campylobacter

Real-time PCR was performed with forward primer 5’-CTGCTTAACACAAGTTGAGTAGG-3’ and reverse primer TTCCTTAGGTACCGTCAGAA as combined and optimized by Lübeck *et al* (xx) with LNA probe 5’-6FAM CA[T]CC[T]CCACGCGGCG[T]TGC BHQ1-3’ [50–53]. Shortly, 500 nM primerers and 300 nM probe were added to Brilliant III Ultra-Fast QPCR mastermix (Agilent) or Tth DNA polymerase mastermix (containing: 1x PCR buffer for Tth DNA polymerase, 25 mM MgCl2, 7% glycerol, 0.6 mM dNTPs, 0,2 mg BSA and 1 U Tth polymerase). For the Brilliant mastermix was the cycling conditions were 95° degrees for 3 minutes followed by 45 cycles of 95° degrees for 5 seconds and 60° degrees for 15 seconds. For the Tth DNA polymerase mastermix was the cycling conditions were 95° degrees for 3 minutes followed by 45 cycles of 95° degrees for 15 seconds, 60° degrees for 60 seconds and 72° degrees for 30 seconds. In both cases a CFX96 instrument (Bio-Rrad) was used.

#### Metagenomic library preparation & metagenomics shotgun sequencing

Metagenomic shotgun libraries were prepared at the Norwegian Sequencing Centre (Oslo, Norway) from the air filter DNA extracts and the two *C. jejuni* isolates. DNA was mechanically fragmented using sonication to 350 bp (Covaris) and Dual indexed libraries were generated using SMARTer ThruPLEX (Takara) adapted to low-input DNA. The libraries were sequenced on a HiSeq 3000 (Illumina) generating 150 bp Paired end reads.

#### Genome assembly of Campylobacter genomes

The *Campylobacter* sequence data was processed with the Bifrost pipeline [54] (https://github.com/NorwegianVeterinaryInstitute/Bifrost) with modifications. In brief, PhiX sequences were removed using BBmap (v 38.76) [55], adapter sequences and low quality bases were removed using Trimmomatic (v 0.39) [56]. Assembly was performed with Skesa (v 2.3.0) [57], using default settings. Due to the very high coverage (> 2000 X, we screened the contigs for contamination using Kraken2 (v 2.0.8_beta) [58] and a Minikraken database containing Refseq microbial genomes. We identified a contaminant *Staphylococcus epidermis* genome (accession number: CAJPVF01) in the *C. jejuni 927* assembly that was present as ≈ 1% of the reads. All non-Campylobacter contigs were removed from the *C. jejuni* 927 assembly. Next, we use Pilon (v 1.23) [59] to correct the assemblies of both isolates and Prokka (v 1.14.5) [60] was used to annotate both genomes.

#### Metagenomic data quality control and classification

Metagenome sequence data quality control was performed using the Sunbeam pipeline (v 2.1.0) [61] (Supplementary materials figure S4). In brief, the pipeline runs Cutadapt (v 2.8) and Trimmomatic (v 0.36) to remove adapters and low quality bases. Next, low-complexity regions are masked using Komplexity (v 0.3.6) and subsequently host contamination filtering was performed to remove read matching both the PhiX (NC_001422.1) and a masked version of the human genome (Hg19, https://drive.google.com/file/d/0B3llHR93L14wd0pSSnFULUlhcUk/edit?usp=sharing). The sequence depth for the individual samples was estimated using Nonpareil (version3) [26]. Taxonomic classification was performed using Kraken (v 1.0) [27]. Species abundances were subsequently estimated using Bracken (v2.6.0) [62].

#### Mapping of metagenome sequences

Species abundances based on reads were also determine in the following way: a collection of fasta sequences was downloaded that included the mock community species genomes (https://s3.amazonaws.com/zymo-files/BioPool/ZymoBIOMICS.STD.refseq.v2.zip), 29 Complete *Campylobacter* spp. reference genomes from Refseq (downloaded 25-11-2020, Supplementary table S1), and the *Gallus gallus* reference genome version 6 (GCA_000002315.5). We also added the genome of *C. coli* NTICC13 (GCF_009756375.1). All genomes were masked using BBmask (BBTools version 38.86), with entropy set to 0.7 [55]. Metagenomic reads were mapped against the masked genomes using BBsplit with default settings and discarding reads with ambigious mapping within a genome or between genomes. A summary of the mapped reads per genome was used for visualization.

#### Visualization of the results

The figures were created using R-studio (version 1.3.959) with R (version 4.0.0). The following R-packages were used to process data tables and generate figures: dplyr (v1.0.2), forcats (v 0.5.0), ggplot2 (v3.3.2), purr (v 0.3.4), readr (v1.4.0), tidyr (v1.1.2).

## List of abbreviations

## Declarations

### Ethics declarations

“Not applicable”

### Availability of data and Materials

All sequence data was submitted to the ENA database and is publically available under the umbrella project PRJEB44024. The individual project numbers are: Genome assemblies - PRJEB43619; Genome fastq files - PRJEB43621; Metagenome fastq files - PRJEB43623. For sample specific accession numbers, please see Supplementary materials table S1 and S3.

### Authors’ contributions

Conception and design: GJ,CS,MT. Drafting of manuscript: TH, CS, GJ, BS. Review and revision of manuscript: TH, CS, GJ, BS, MT. Sample collection: CS, GJ, MT. Analysis and data interpretation: TH, CS, GJ, BS. The authors read and approved the final manuscript.

## Acknowledgements

We thank the Norwegian Sequencing Centre for support, library preparation and sequencing of our samples. The analysis of the metagenomics data was performed under the project code NN9305K on 1) the Abel Cluster, which is operated by the Department for Research Computing at USIT and owned by the University of Oslo and Uninett/Sigma2, and 2) the Saga Cluster, owned and maintained by Uninett/Sigma2. (Sigma2.no). The farmer association and farmer that provided access to poultry houses are acknowledged for their kind collaboration. Tone Fagereng Mathisen, NVI is acknowledged for assisting with the *Campylobacter* isolates used in the spiking experiments.

## Electronic supplementary material

Supplementary materials.pdf

