## Supplementary Materials for "Detection of *Campylobacter* in air samples from poultry houses using shot-gun metagenomics – a pilot study"

### Tables

**Supplementary materials table S1**. Genome characteristics and accession numbers of the *Campylobacter jejuni* isolates used in this study.

| **Strain** | **Illumina PE reads** | **No. of contigs** | **Total size (Mbp)** | **GC content (%)** | **No. of genes** | **Average genome coverage** | **Reads accession no.** | **Genome assembly accession no.** |
|---|---|---|---|---|---|---|---|---|
| 927 | 19662413 | 32 | 1.59 | 30.4 | 1622 | 356 | ERS6103276 | CAJPVE01 |
| CCUG11284T | 20087479 | 53 | 1.73 | 30.2 | 1831 | 2837 | ERS6103275 | CAJQFQ01 |

**Supplementary materials table S2**. Overview of complete *Campylobacter* reference genomes used for mapping. Selected and downloaded on 25-11-2020.

| Species | Strain | Assembly accession | Size (Mb) | GC content (%) | Isolation source |
|---|---|---|---|---|---|
| *Campylobacter rectus* | ATCC 33238 | GCA_004803795.1 | 2,57 | 44,7 | human peridontal pocket |
| *Campylobacter showae* | B91_SC | GCA_900699785.1 | 2,16 | 46,0 | human oral cavity |
| *Campylobacter gracilis* | ATCC 33236 | GCA_001190745.1 | 2,28 | 46,6 | human gingival sulcus |
| *Campylobacter insulaenigrae* NCTC 12927 | NCTC 12927 | GCA_000816185.1 | 1,47 | 28,2 | marine mammal. |
| *Campylobacter avium* LMG 24591 | LMG 24591 | GCA_002238335.1 | 1,74 | 34,2 | poultry |
| *Campylobacter lanienae* NCTC 13004 | NCTC 13004 | GCA_002139935.1 | 1,59 | 34,6 | Human feacal sample |
| *Campylobacter mucosalis* | ATCC 43264 | GCA_013372205.1 | 1,77 | 36,6 | porcine small intestine |
| *Campylobacter volucris* LMG 24379 | LMG 24379 | GCA_000816345.1 | 1,52 | 28,6 | rectal swab black-headed gulls |
| *Campylobacter subantarcticus* LMG 24377 | LMG 24377 | GCA_000816305.1 | 1,85 | 29,8 | Gray-headed albatros |
| *Campylobacter corcagiensis* | LMG 27932 | GCA_013201645.1 | 1,69 | 31,9 | lion-tailed macaques feaces |
| *Campylobacter hepaticus* | HV10 | GCA_001687475.2 | 1,52 | 28,0 | Chicken liver with spotty liver disease |
| *Campylobacter ornithocola* | LMG 29815 | GCA_013201605.1 | 1,64 | 29,2 | Wild bird feacal samples |
| *Campylobacter geochelonis* | LMG 29375 | GCA_013201685.1 | 2,17 | 33,5 | western Hermann's tortoise |
| *Campylobacter blaseri* | LMG 30333 | GCA_013201895.1 | 1,89 | 29,3 | Common seals |
| *Campylobacter canadensis* | LMG 24001 | GCA_013177655.1 | 1,92 | 27,4 | captive whooping cranes |
| *Campylobacter fetus subsp. testudinum* Sp3 | SP3 | GCA_001484645.1 | 1,82 | 33,1 | Humans and reptiles |
| *Campylobacter lari* RM2100 | RM2100 ATCC BAA-1060D | GCA_000019205.1 | 1,57 | 29,6 | Seabirds and mammals |
| *Campylobacter upsaliensis* RM3940 | RM3940 | GCA_013372245.1 | 1,63 | 35,1 | Humans and mammals |
| *Campylobacter curvus* 525.92 | 525.92 | GCA_000017465.2 | 1,97 | 44,5 | Human |
| *Campylobacter helveticus* | ATCC 51209 | GCA_002080395.1 | 1,87 | 34,3 | mammals |
| *Campylobacter cuniculorum* DSM 23162 = LMG 24588 | LMG 24588 | GCA_002104335.1 | 1,94 | 31,2 | Rabbit ceaceum |
| *Campylobacter hyointestinalis subsp. lawsonii* | CHY5 | GCA_013372165.1 | 1,81 | 33,3 | porcine stomach |
| *Campylobacter sputorum bv. Paraureolyticus* LMG 11764 | LMG 17589 | GCA_002220755.1 | 1,73 | 29,6 | human |
| *Campylobacter ureolyticus* RIGS 9880 | RIGS 9880 | GCA_001190755.1 | 1,64 | 29,2 | Human and mammals |
| *Campylobacter peloridis* LMG 23910 | LMG 23910 | GCA_000816785.1 | 1,76 | 28,4 | shellfish |
| *Campylobacter iguaniorum* | 1485E | GCA_000736415.1 | 1,75 | 35,8 | reptiles |
| *Campylobacter pinnipediorum subsp. caledonicus* | RM18020 | GCA_002021985.1 | 1,71 | 30,4 | Seals |
| *Campylobacter armoricus* | CCUG 73571 | GCA_013372105.1 | 1,64 | 28,6 | Marine samples and human stool |
| *Campylobacter jejuni subsp. jejuni* NCTC 11168 = ATCC 700819 | NCTC 11168 | GCA_000009085.1 | 1,64 | 30,5 | Human feces |
| *Campylobacter coli* | NTICC13 | GCF_009756375 | 1,74 | 31,3 | unknown |

**Supplementary materials table S3**. Overview of shotgun metagenome accession numbers as used in the ENA database.

| Samples | NCBI Taxon id | Study accession | Sample accession | Experiment accession | Run accession |
|---|---|---|---|---|---|
| 1-MOCK1-B-a | 1235509 | PRJEB43623 | ERS6130789 | ERX5340133 | ERR5621728 |
| 2-MOCK1-B-b | 1235509 | PRJEB43623 | ERS6130790 | ERX5340134 | ERR5621729 |
| 3-MOCK1-S1-a | 1235509 | PRJEB43623 | ERS6130791 | ERX5340135 | ERR5621730 |
| 4-MOCK1-S1-b | 1235509 | PRJEB43623 | ERS6130792 | ERX5340136 | ERR5621731 |
| 5-MOCK1-S2-a | 1235509 | PRJEB43623 | ERS6130793 | ERX5340137 | ERR5621732 |
| 6-MOCK1-S2-b | 1235509 | PRJEB43623 | ERS6130794 | ERX5340138 | ERR5621733 |
| 7-MOCK2-B-a | 1235509 | PRJEB43623 | ERS6130795 | ERX5340139 | ERR5621734 |
| 8-MOCK2-B-b | 1235509 | PRJEB43623 | ERS6130796 | ERX5340140 | ERR5621735 |
| 9-H1-3 | 655179 | PRJEB43623 | ERS6130797 | ERX5340141 | ERR5621736 |
| 10-H1-4 | 655179 | PRJEB43623 | ERS6130798 | ERX5340142 | ERR5621737 |
| 11-H1-5 | 655179 | PRJEB43623 | ERS6130799 | ERX5340143 | ERR5621738 |
| 12-H1-B | 655179 | PRJEB43623 | ERS6130800 | ERX5340144 | ERR5621739 |
| 13-H2-3 | 655179 | PRJEB43623 | ERS6130801 | ERX5340145 | ERR5621740 |
| 14-H2-4 | 655179 | PRJEB43623 | ERS6130802 | ERX5340146 | ERR5621741 |
| 15-H2-5 | 655179 | PRJEB43623 | ERS6130803 | ERX5340147 | ERR5621742 |
| 16-H2-B | 655179 | PRJEB43623 | ERS6130804 | ERX5340148 | ERR5621743 |

### Figures

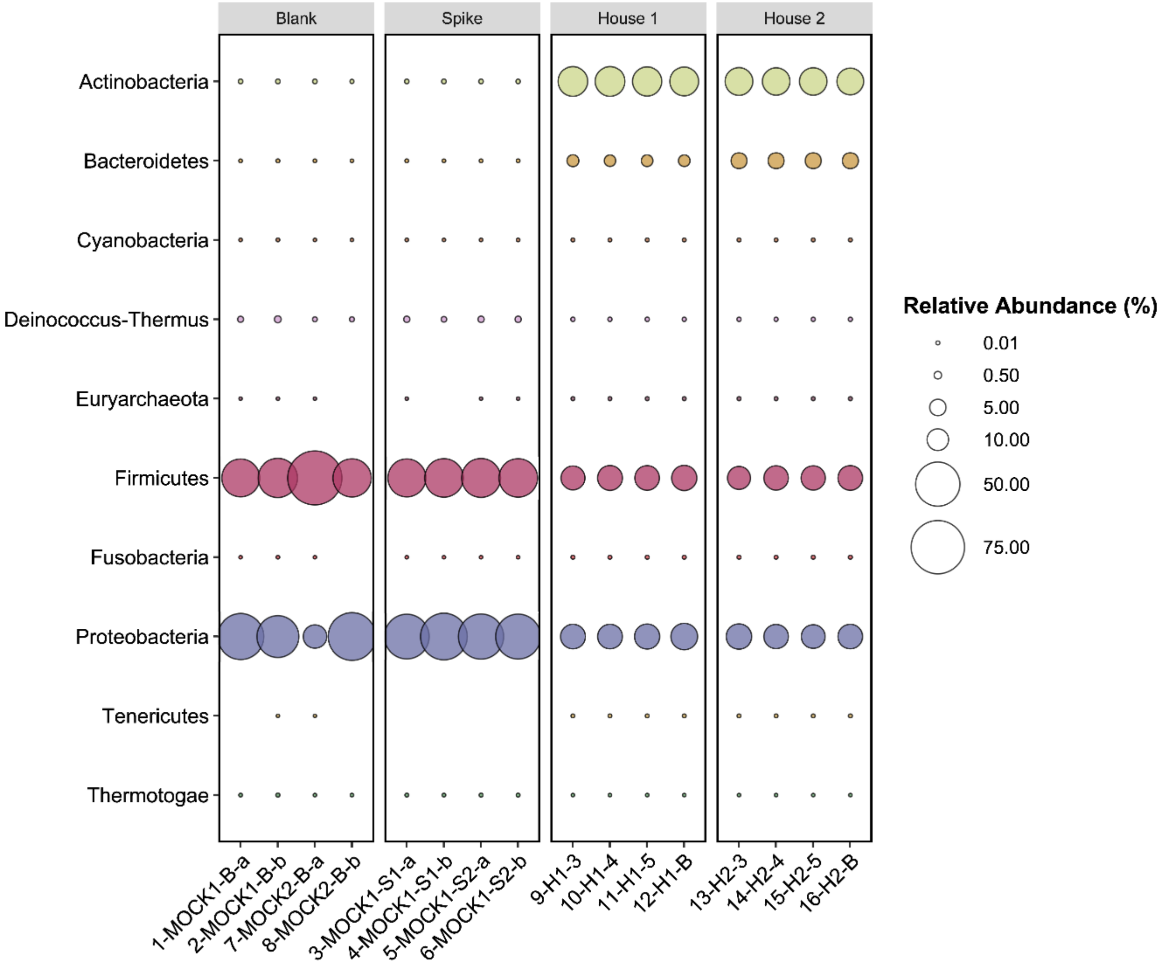


**Supplementary materials figure S1**: Relative abundances of bacterial phyla found in MOCK and HOUSE samples. Phyla with a relative abundance equal or higher than 0.01% in at least one sample were kept for visualization. Note, relative abundances below 0.01% are not shown.

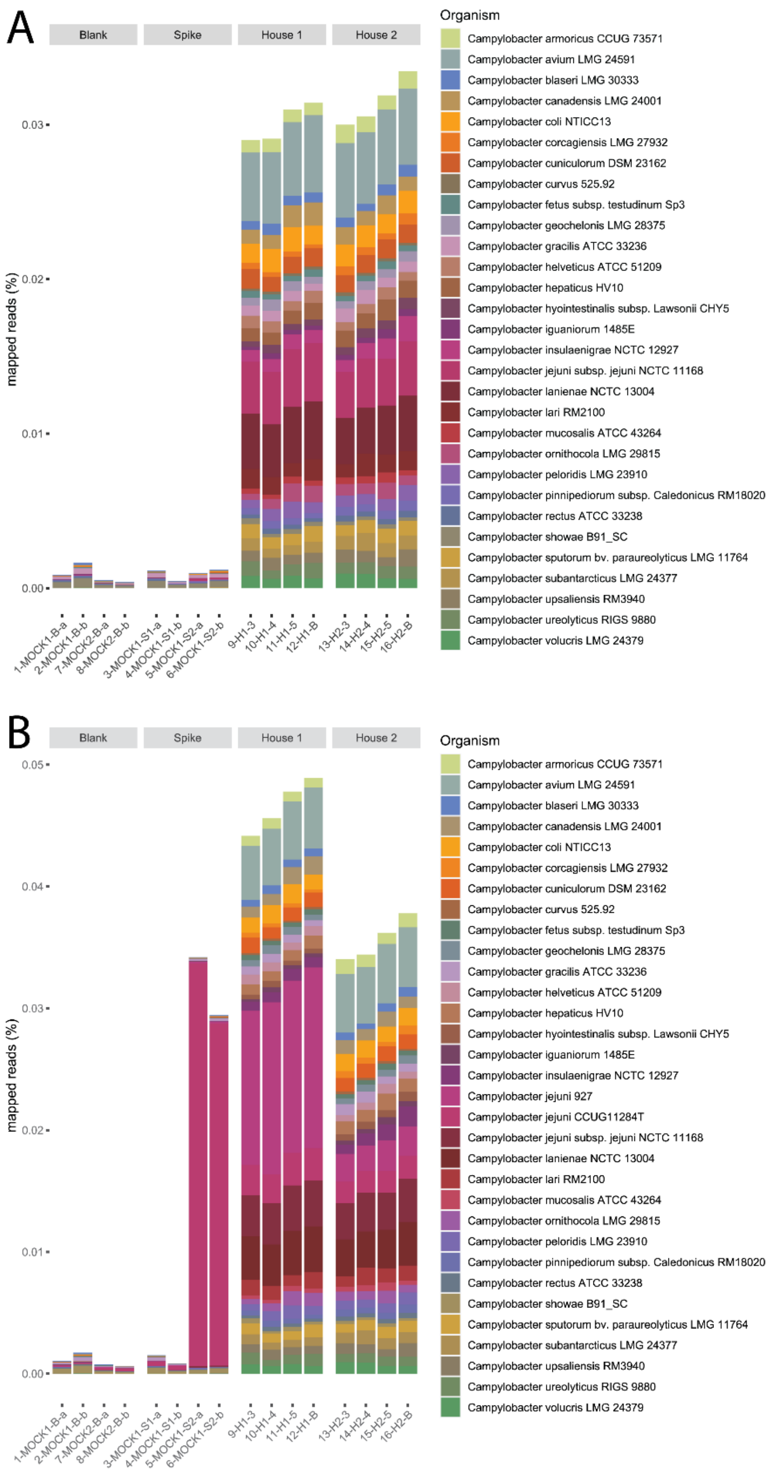


**Supplementary materials figure S2**: Relative abundances of *Campylobacter* spp. in MOCK and HOUSE samples. Metagenomic reads were mapped with BBsplit to a selection of *Campylobacter* spp. reference genomes (A), or to the same selection including the genomes from our spiked in isolates: *C. jejuni* CCUG 11284T (MOCK samples) and *C. jejuni* 927 (HOUSE samples).

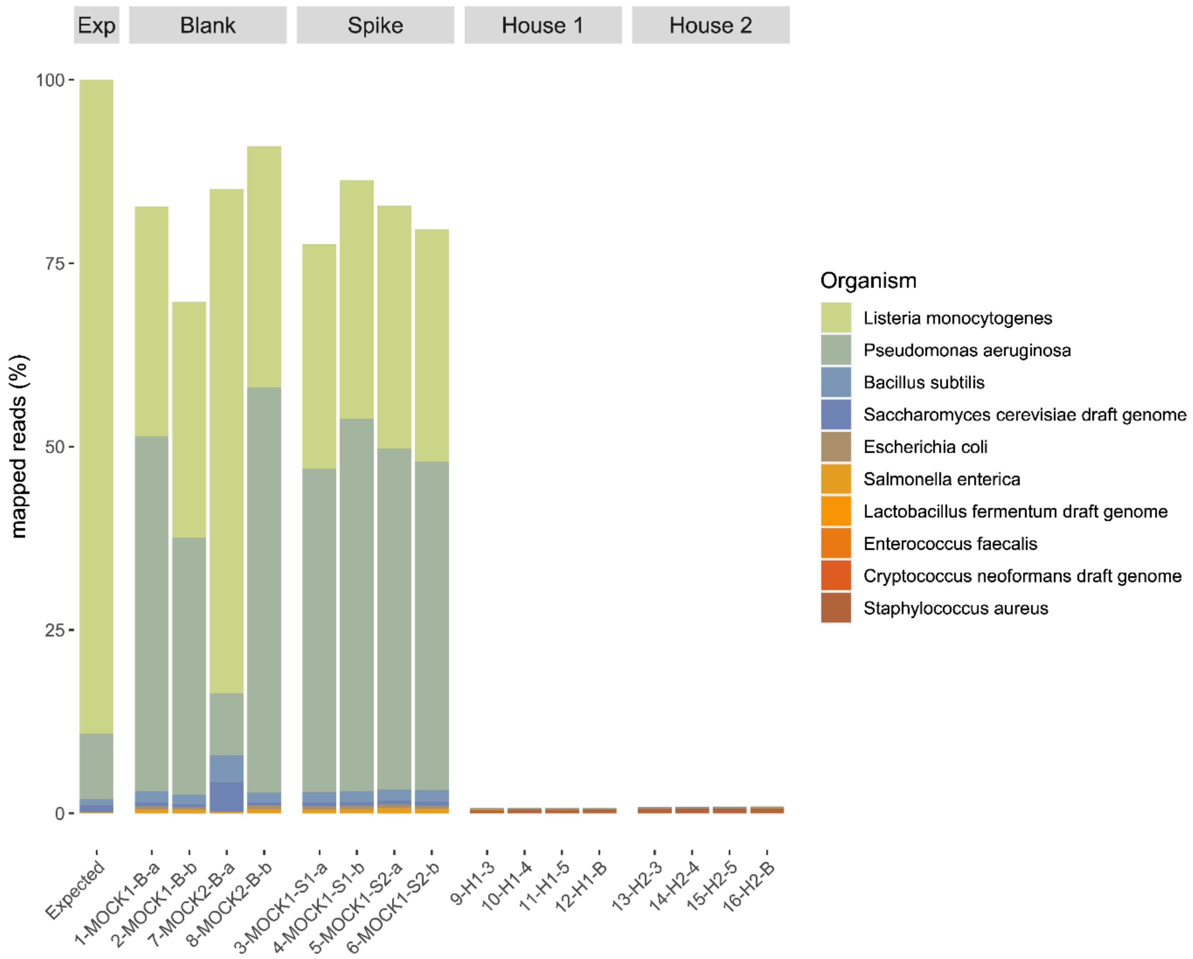


**Supplementary materials figure S3**: Relative abundances of mock community taxa in MOCK and HOUSE samples. Metagenomic reads were mapped with BBsplit to the genomes of the isolates used in the construction of the Mock community. The expected relative abundances of the taxa are: *Listeria monocytogenes*: 89.1% ; *Pseudomonas aeruginosa*: 8.9% ; *Bacillus subtilis*: 0.89% ; *Saccharomyces cerevisiae*: 0.89% ; *Escherichia coli*: 0.089% ; *Salmonella enterica*: 0.089% ; *Lactobacillus fermentum*: 0.0089% ; *Enterococcus faecalis*: 0.00089% ; *Cryptococcus neoformans*: 0.00089% ; *Staphylococcus aureus*: 0.000089%.

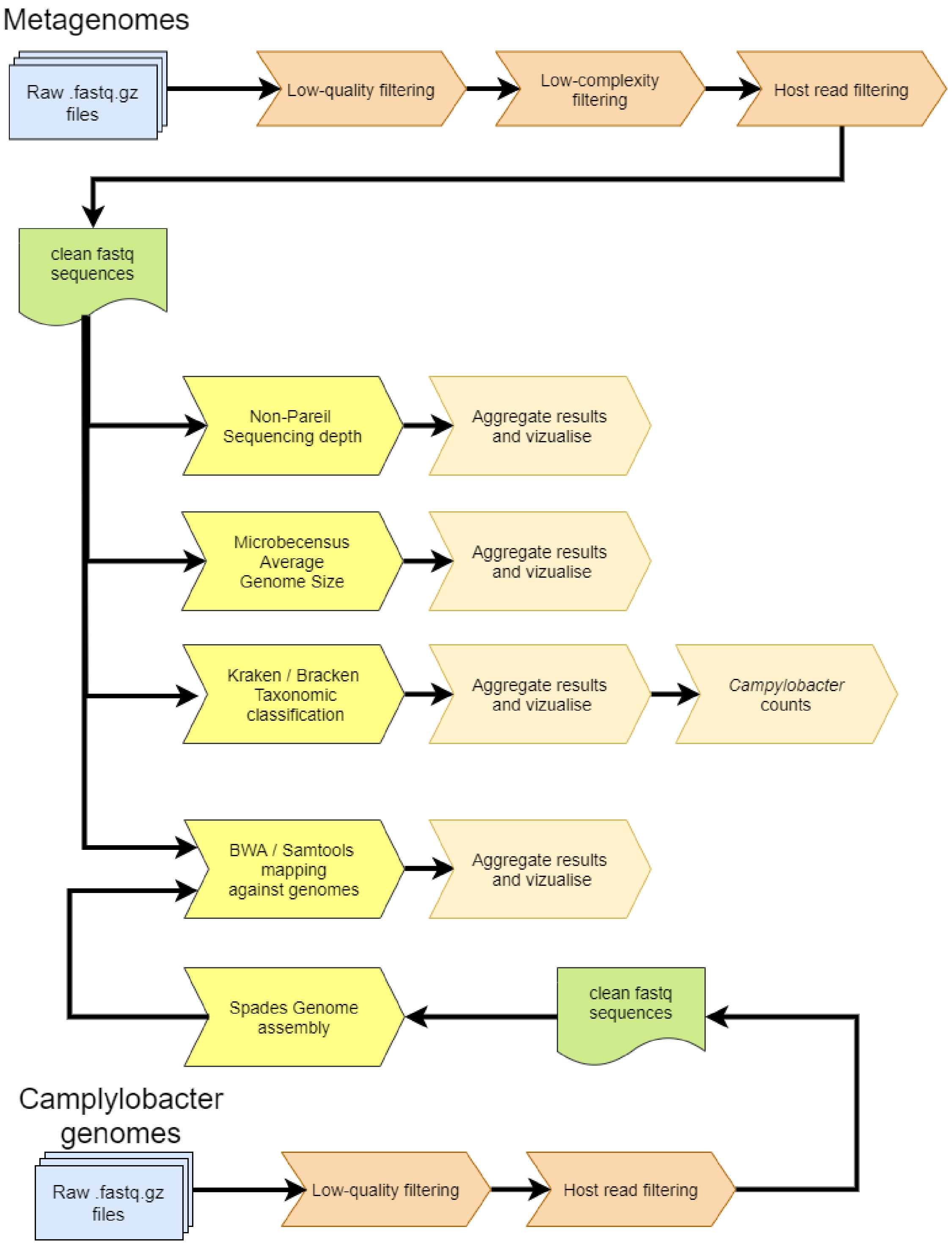


**Supplementary materials figure S4**: Schematic overview of the bioinformatics workflow used in this project.
